# RB-TnSeq analysis reveals alcohol and salt tolerance systems in a plant root colonizer *Paraburkholderia graminis* OAS925

**DOI:** 10.64898/2026.01.28.702333

**Authors:** Shweta Priya, Thomas Eng, Valentine Trotter, Adam M. Deutschbauer, Jenny C. Mortimer, Aindrila Mukhopadhyay

**Affiliations:** Biological Systems and Engineering Division, Lawrence Berkeley National Laboratory, Berkeley, CA, USA; Environmental Genomics & Systems Biology Division, Lawrence Berkeley National Laboratory, Berkeley CA, USA; Department of Plant and Microbial Biology, University of California, Berkeley, CA, USA; School of Agriculture, Food and Wine & Waite Research Institute, Adelaide University, Adelaide, South Australia 5005, Australia

## Abstract

The role of microbial strains in regulating natural stresses and their impact on plant health is well-established. However, the role of microbial tolerance mechanisms in plant response to unnatural or anthropogenic stresses is less understood. Examination of these interactions impact our deeper understanding of plant-microbe interactions and our ability to enhance beneficial functions. In this study we use the model plant *Brachypodium distachyon* and its prominent root colonizer *Paraburkholderia graminis* OAS925 to investigate mechanisms of tolerance to alcohol and salt stress. We examined the ability of OAS925 to reduce root growth inhibition during exposure to short chain alcohols and salt. We also examined the tolerance mechanism for OAS925 towards these stresses using RB-TnSeq fitness assays. The most prominent tolerance systems in OAS925 are genes specifically involved in membrane transport (such as the Mla operon), efflux systems (e.g., RND efflux systems), signaling and regulation (PrtR/PrtI, NtrY/NtrX, and EnvZ/OmpR), and oxidative stress response (GshB). Our findings provide a model where bacterial membrane integrity, active solvent efflux, and stress signaling are crucial not only for bacterial survival but also for maintaining the root colonization and biofilm formation that confer protection to the host plant.

## 1. Introduction

Advancements in biomanufacturing are leading to the development of new biorefineries, and targeting many compounds, including fuel precursors and platform chemicals. A critical area of investigation is the evaluation of the environmental consequences associated with these newly introduced chemicals especially in the context of crop health, plant growth, and overall yield [1]. Reports in the literature show that the impact of these solvents on the plants include damage to cell membranes caused by oxidative stress, decreased gas exchange, inhibited seed germination, and potential plant death at higher concentrations [2].

The beneficial role of the plant-associated microbiome in promoting rhizosphere health and plant vigor is well established. Numerous studies have highlighted the importance of plant-microbe interactions in protecting plants from both biotic and abiotic stresses [3,4]. However, while benefits of the associated plant microbiome are extensively documented, impact of the exogenously introduced novel anthropogenic chemicals on these microbes and their ability to protect the host plants is less understood. Additionally, despite the known high tolerance of plant-associated microbes to environmental perturbations, most research on bacterial mechanisms for alcohol toxicity tolerance has concentrated on model bacteria, which may not accurately represent the behavior of rhizobial strains and in plant-associated environments [5,6].

*Paraburkholderia graminis* OAS925 (previously known as *Burkholderia* OAS925 and referred to here on as OAS925) is an efficient root colonizer of *Brachypodium distachyon* and known for its high stress tolerance and plant growth benefits [7–9]. It is a dominant and well-established root colonizer across many *Poaceae* members (grasses) and is supported by robust functional genomic tools [9–11]. As such, OAS925 provides an ideal system to study microbial tolerance mechanisms and corresponding ability to protect the host *B. distachyon,* from various chemical toxicities.

We investigated a selection of short-chain alcohols and salt, chosen as representative bioderivable anthropogenic chemicals. Isoprenol and Prenol, collectively referred to as isopentenols, is an increasingly sought-after industrial solvents due to their ability to serve as precursors for synthesizing various valuable chemicals including advanced biofuels [12,13]. Despite the increasing demand and intensive research to improve the industrial production of isopentenols, their ecological impacts, particularly within plant environments, are still unexplored. To determine if alcohols with similar chemical structures exhibit comparable toxicity to plant-associated microbiota, we also included ethanol, 2-propanol and 1-propanol in our study. Salt was chosen as a relevant desiccation-related stressor, as it occurs naturally in plant environments and is also a potential industrial contaminant.

We evaluated the impact of these stressors on microbial response and in plant systems, via extensive functional genomics. Specifically, we use a random barcode transposon-site sequencing (RB-TnSeq) mutant library of OAS925, to reveal the major genetic determinants in conferring bacterial tolerance to these short chain alcohols. We also characterize the impact of OAS925 root inoculation on plants grown under these inhibitory conditions and evaluate the relationship between bacterial fitness and plant colonization which play a major role in plant’s tolerance to these toxic solvents. In this study we develop both a microbial response framework and the role of microbes in the ecosystem to shield plants from these anthropogenically introduced chemical stresses.

## 2. Materials and Methods

### 2.1 Microbial growth conditions and alcohol sensitivity assays

Five short chain alcohols 3-methyl-3-buten-1-ol (isoprenol), 3-methyl-2-buten-1-ol (prenol), ethanol, 2-propanol, and 1-propanol; and salt (NaCl) were used to assay response in the native rhizosphere bacteria OAS925 (Fig. 1B). All reagents were purchased from Sigma Aldrich (St. Louis, MO, USA) unless otherwise stated. In order to use the OAS925 RB-TnSeq mutant library for functional characterization, first we determined the minimum inhibitory concentration (MIC) for each compound in WT OAS925. In the case of isoprenol and prenol, we used initial alcohol concentrations of 0, 2, 4, 6, 8 and 10 g/L and for 2-propanol and 1-propanol, we used 0, 2, 4, 6, 8, 10, 12, 14, 16, 18 and 20 g/L diluted in liquid M9-glucose medium. For ethanol, we used 0, 2%, 4%, 6% and 8% v/v of liquid M9-glucose medium. For quantification, six replicates per condition were performed. Five mL of bacterial cultures were grown at 200 rpm at 30 °C for 24 hrs. Subsequently, the cultures were centrifuged and inoculated into 24-deep-well plates in a total volume of 3 mL, with an initial inoculum concentration of 0.1 at OD_600_. Plates were sealed with Breathe-Easy® membranes (Sigma-Aldrich, USA Scientific) and were incubated under the same conditions for 24 hrs. The final OD_600_ was quantified using the Spectrophotometer (Molecular Devices, San Jose, CA, USA).

**Fig 1.**
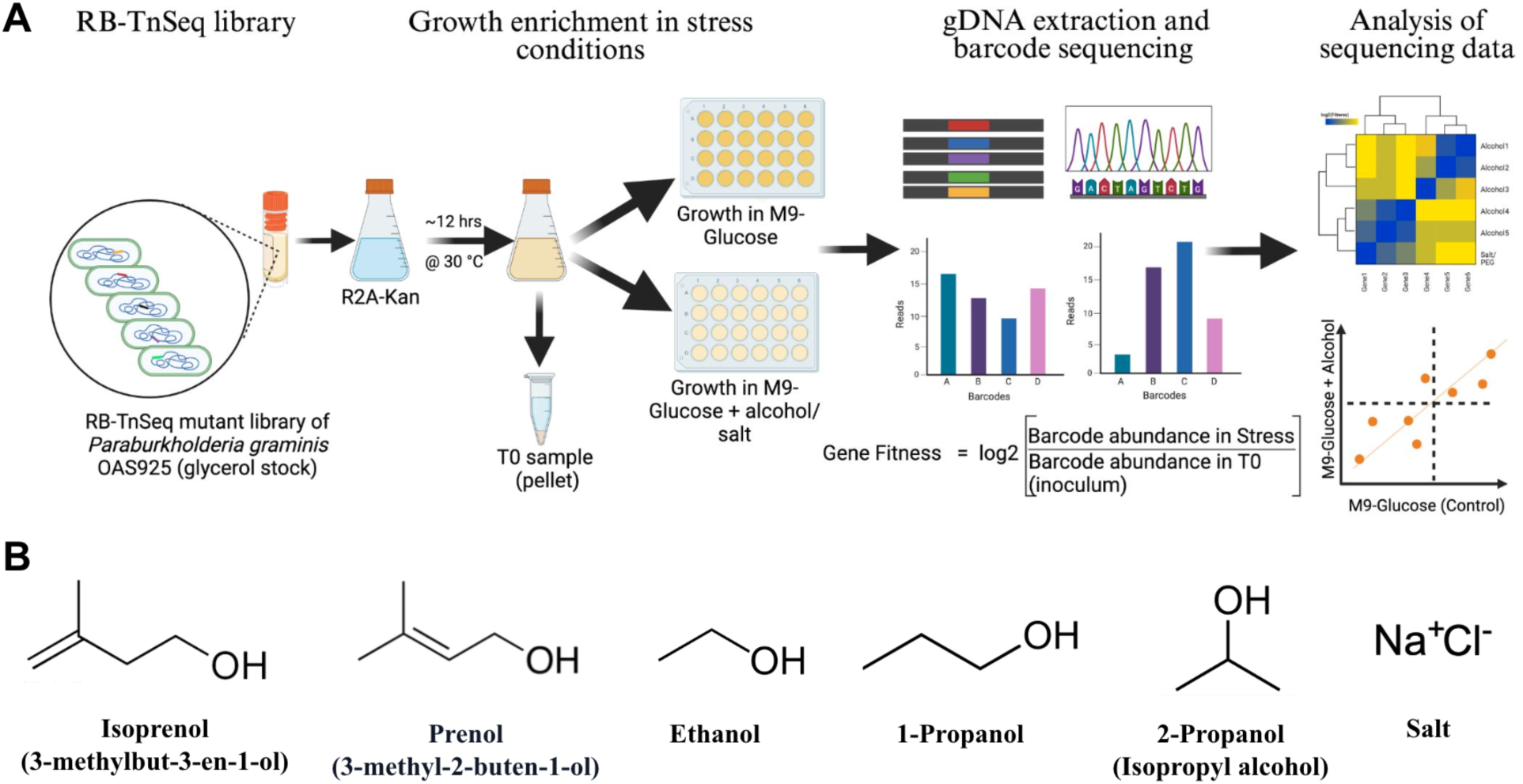
Workflow of the RB-TnSeq fitness profiling for *P. graminis* OAS925 grown under various solvent and salt stress (A). Names and chemical structures of solvents used in the study to test the tolerance of OAS925 (B).

### 2.2 Plant growth conditions and sensitivity assay to alcohols

The set of short chain alcohols were also tested in *B. distachyon* at different concentrations of the stressors. For this, plant seeds were sterilized using 30 sec of 70% ethanol followed by 5 min of 6% bleach and rinsed thoroughly with sterile water, and then germinated on 0.5x Murashike and Skoog (MS) agar medium (Phyto Technology Laboratories, Lenexa, KS, USA) plates. Four days post germination, seedlings were transferred to 0.5 x MS medium plates supplemented with different concentrations (same range as described for microbial MIC analysis above) of alcohols. The various alcohols (see concentration in prior section) were added to the melted media agar and used to pour plates. The seedlings were lifted off of 0.5 x MS media plates using sterile forceps and tapped to remove the attached agar before transferring them to the plates containing alcohol/salt in 0.5 x MS medium. Seedlings with visually similar root lengths were selected for transfer. Seedlings with broken roots during the transfer were discarded, with at least 4 seedlings per plate. For the bacterial inoculation, seedlings were treated with the OAS925 as described in the next section.

### 2.3 Plant root inoculation with OAS925

OAS925 was grown in Reasoner’s 2A (R2A) medium (HiMedia Laboratories LLC, Kennett Square, PA, USA) overnight at 30 °C and shaking at 200 rpm. Overnight cultures were centrifuged (2000 rcf) for 10 min at room temperature, pellet was then resuspended in sterile water and diluted to the OD_600_ of ∼ 0.1. Plants were inoculated by submerging roots into cell resuspensions for approximately 5 min or in sterile water for uninoculated controls. Plants were transferred to fresh 0.5x MS agar plates immediately postinoculation and grown as before for 10 days. Primary root length was then measured using imageJ Fiji software [14].

### 2.4 RB-TnSeq competitive growth assays

To carry out the RB-TnSeq growth assays in this work, we used the previously generated RB-TnSeq library from OAS925 [11]. Briefly, the RB-TnSeq mutant library of OAS925 was generated via conjugation with an *E. coli* WM3064 donor carrying the pHLL250 mariner transposon vector library (strain AMD290), as described in details in [11] and the barcodes were mapped as described in [15].

The experimental workflow for the RBTnSeq growth assays is described in Figure 1A. Briefly, one milliliter aliquots of the frozen stock of the OAS925 mutant library, namely *Paraburkholderia*_OAS925_ML2, were diluted in 25 mL of R2A supplemented with kanamycin (10 μg/mL) and grown for ∼12 hours until the culture reached OD_600_ of 1.0. To collect Time0 (T0) samples, 1mL aliquot from the recovered library was centrifuged and pellet was stored at -20 °C until DNA extraction. The remaining cells were then washed twice with M9 without any carbon source. The washed cells were then grown in either M9 medium containing glucose (mock control) or supplemented with 52 mM isoprenol, 23 mM of prenol, 428 mM of ethanol, 349 mM of 2-propanol, 216 mM of 1-propanol or 200 mM of salt in different 24 deep well plates. These concentrations reflect the experimentally determined MIC for each alcohol. For each experiment, the mutant library was inoculated to a starting OD_600_ of 0.02. Four biological replicates were performed for each condition. Subsequently, the bacterial libraries were cultivated at 30 °C with shaking at 200 rpm for ∼24 hours. Well contents were transferred to an 1.7 ml eppendorf tube and centrifuged at 6010 x*g* for 3 min, and the supernatant was discarded. Mutant library cell pellets were stored at -20 °C until genomic DNA extraction. Genomic DNA of each RB-TnSeq sample that grew in uninoculated media or in the media with the stressors were extracted using Quick-DNA Fungal/Bacterial Miniprep Kit from Zymo Research. Subsequently, PCR was performed to amplify only the barcodes in each sample using universal primers, and amplicons were sequenced on an Illumina NovaSeq instrument, as described [15].

### 2.5 RB-TnSeq gene fitness calculation

The fitness of a strain is the normalized log2 ratio of barcode reads in the experimental sample to barcode reads in the time zero sample. The fitness of a gene is defined here as the weighted average of the strain fitness for insertions in the central 10% to 90% of the gene. The gene fitness values are normalized so that the typical gene has a fitness of zero. The primary statistical *t* value represents the form of fitness divided by the estimated variance across different mutants of the same gene. Statistical |*t*| of > 4 were considered significant. A fully detailed explanation of calculating fitness scores are reported in Wetmore et al. [15]. All the experiments described here passed testing using the quality metrics described previously unless otherwise noted [15]. All experiments were conducted with 4 biological replicates and all fitness data are available at http://fit.genomics.lbl.gov.

Comparison of mean fitness values between the alcohol/salt added medium and medium reference cultures (M9 + 20 mM glucose alone) was performed using a two-sample *t*-test, where the *p* value was corrected for multiple testing via the positive false discovery rate (pFDR) method [16].

### 2.6 Growth test with insertional mutants in single genes

To validate fitness results obtained using OAS925 RB-TnSeq mutant pools, we performed additional growth assays with individual mutants. Single transposon mutants of *Paraburkholderia_*OAS925_ML2, were obtained from an arrayed collection of individual transposon mutants as described in detail in [11]. The single mutant strains from glycerol stocks of the arrayed collection were then revived by streaking on R2A agar and subsequently subcultured in R2A broth followed by incubation at 30 °C and shaking at 200 rpm.

The media used for growth and subculturing of mutant strains was supplemented with kanamycin (10 µg mL^-1^). These mutants are listed in Table 1.

**Table 1.**
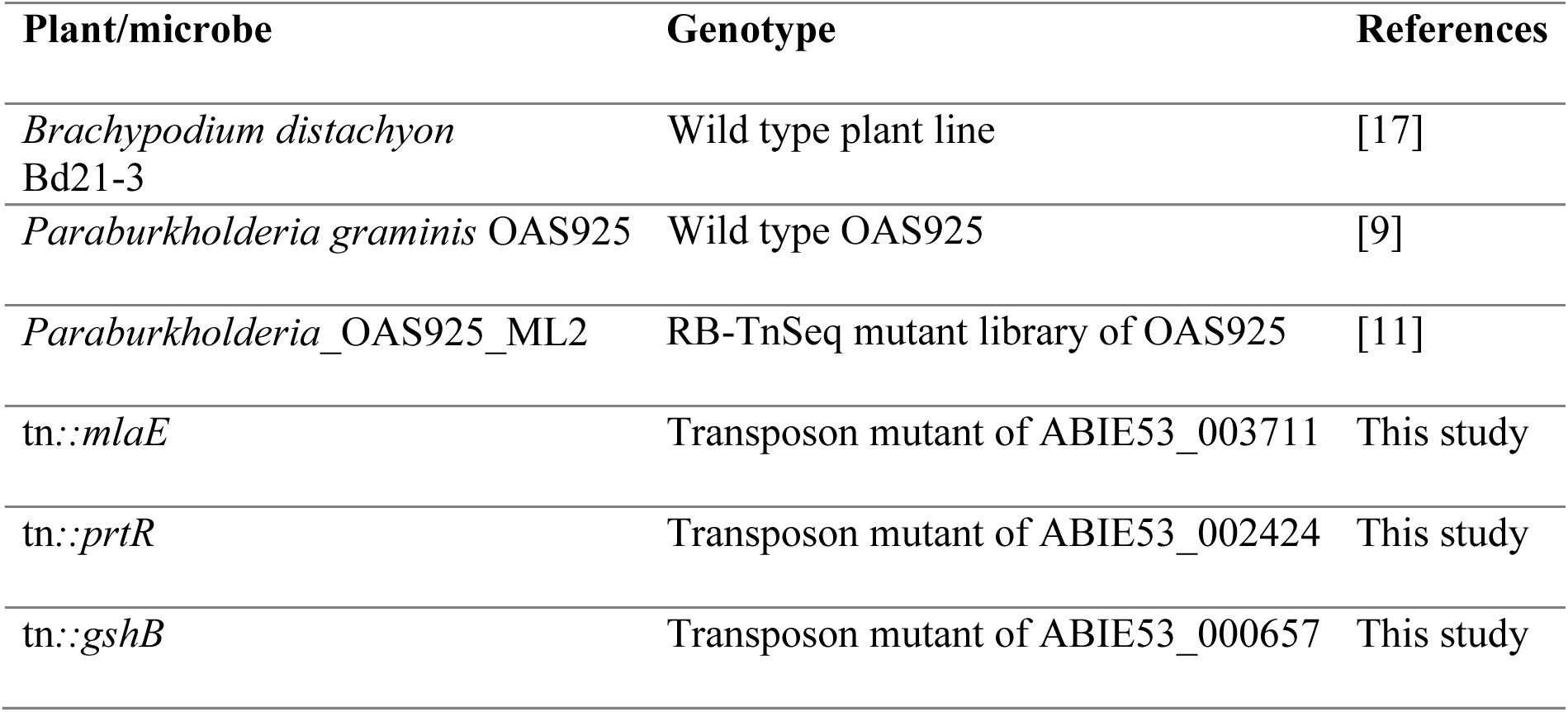
List of plant lines and microbial strains used in the study.

The single mutants were then tested with the alcohols/salt stressors to validate the RB-TnSeq results obtained previously. Overnight cultures of the OAS925 WT and mutants were centrifuged and pellets were resuspended in fresh M9-medium and diluted to the OD_600_ of ∼0.1. The same amount of stressors were added in 48 well culture plates with a reaction volume of 300 μL and OD_600_ was measured every 10 minutes after shaking on a SpectraMax M2e Microplate Reader (Molecular Devices Inc, San Jose, CA, USA).

### 2.7 Plant root colonization and biofilm assay

OAS925 and the mutants (tn*::prtR* and tn*::mlaE*) were grown in Reasoner’s 2A (R2A) medium and R2A supplemented with kanamycin (10 µg mL^-1^) respectively, for overnight at 30 °C and shaking at 200 rpm. Plants were inoculated in the same way as mentioned previously (see section 2.3) and were transferred to fresh 0.5x MS agar plates containing alcohols/salt immediately postinoculation and grown for 10 days.

For colonization assay, roots were immersed in 5mL sterile PBS solution and vortex for ∼10 minutes to separate the cells in the suspension. One mL of this suspension was then pipetted into a microcentrifuge tube, serially diluted and plated on R2A (for WT) and R2A with 10 µg mL^-1^ of kanamycin (for tn*::prtR*) and colonies were counted after 24 hours of incubation at 30 °C. Biofilm was measured using a method described by O’Toole [18] with some modifications. Briefly, inoculated roots were fragmented, and ∼0.5 g placed in sterile 6-well Corning® Falcon® cell culture plates. Three mL of sterile water was added to each well and incubated with shaking (100 rpm) for 30 min to dislodge non-adhering cells. The water was removed, and roots were washed three times with Phosphate-buffered saline (PBS), then air-dried for 15 min. The plates were then submerged in sterile water to remove remaining unattached cells. Roots were stained by adding 250 μL of 0.1% crystal violet solution and incubating for 10–15 min at room temperature. After rinsing the roots 3–4 times with sterile water, 500 μL of 30% acetic acid was added to each well and incubated for 10–15 min at room temperature to solubilize the attached crystal violet. Finally, 250 μL of the solubilized crystal violet was transferred to a new 48 well microtiter plate, and absorbance was measured at 550 nm using a Spectrophotometer (Molecular Devices, San Jose, CA, USA).

### 2.8 Data Analysis

GraphPad Prism 10 (GraphPad Software, San Diego, CA, USA) was used to make the plots and growth curves and perform data analysis. Heatmaps and bubbleplot were created using the ‘seaborn’ package in python [19]. The Principal Component Analysis (PCA) plot was created using the ‘matplotlib’ library in python [20].

## 3. Results and Discussion

### 3.1. Sensitivity of plants to short chain alcohols and salt

To examine the sensitivity of *B. distachyon* (BD21-3) to selected industrial solvents, plants were grown for 2 weeks post germination on 0.5x MS-agar media with ∼0.6 mM isoprenol, 170 mM ethanol, 330 mM 2-propanol and 75 mM salt and measured the root length and number of lateral roots. Additionally, to understand if the addition of OAS925 improves the plant’s alcohol and salt tolerance, plants were inoculated with OAS925 before placing them on media plates prepared with these solvents.

Figure 2A provides representative images (photographs) of *B. distachyon* seedlings after 10 days of growth under control conditions or exposure to isoprenol, ethanol, 2-propanol, or salt, comparing uninoculated plants to those inoculated with OAS925. Quantitative analyses of these phenotypes are presented for primary root length (Fig. 2B) and the number of lateral roots (Fig. 2C). As shown in Figure 2B, exposure to all tested chemical stressors significantly inhibited primary root elongation in uninoculated plants compared to the unstressed control. For example, exposure to 170 mM ethanol reduced primary root length by ∼ 50%, from a mean of 6.83±0.70 cm to 3.56±1.13 cm. The most important observation was that inoculation with OAS925 significantly mitigated this growth inhibition across all stress and no stress conditions. In the ethanol treatment, for instance, inoculation with OAS925 increased the primary root length by ∼1.5X compared to the uninoculated, stressed plants. A similar protective effect was observed for lateral root development under ethanol and salt stress (Fig. 2C). However, lateral root numbers varied with different kinds of compounds under inoculated vs uninoculated conditions.

**Fig 2.**
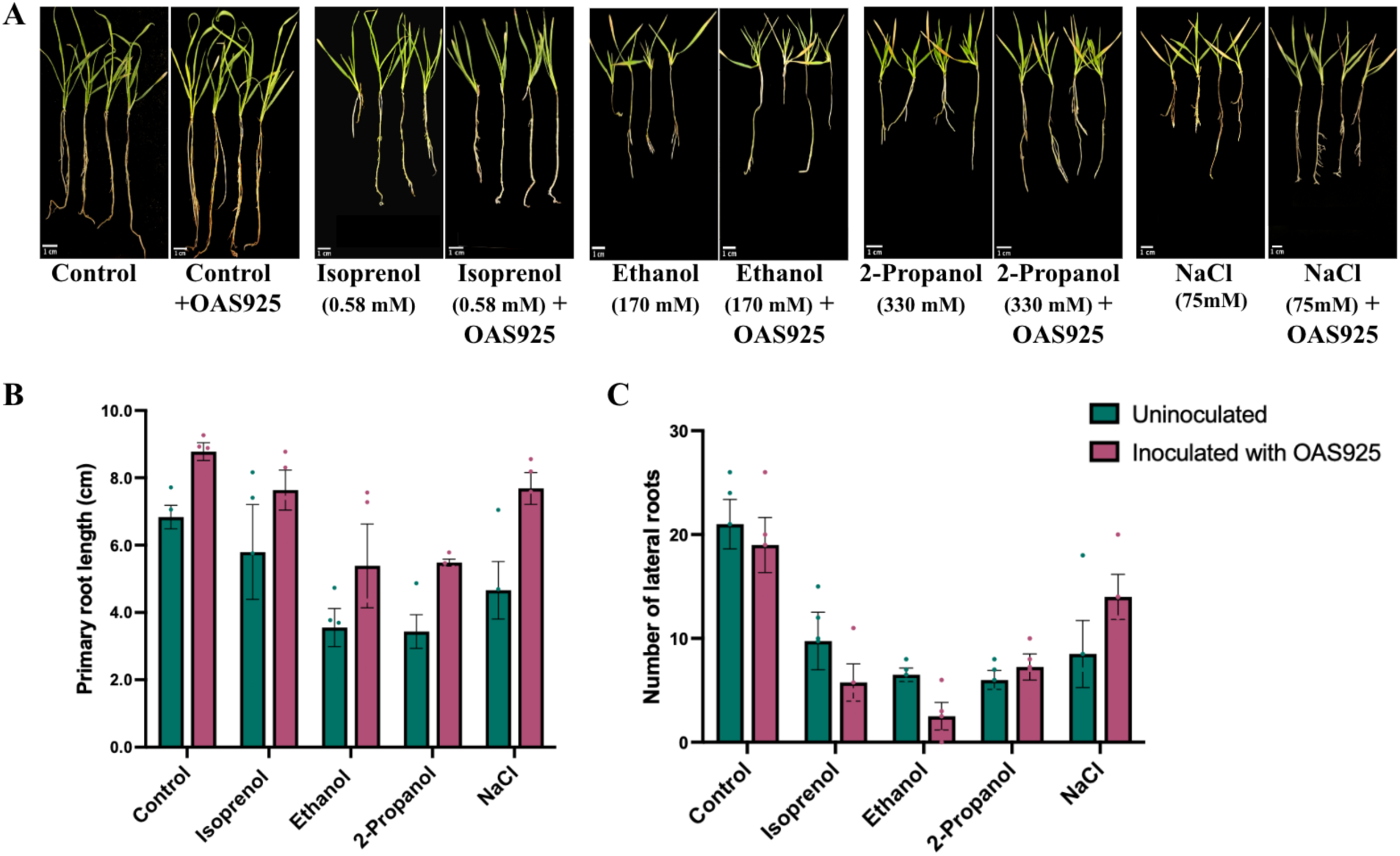
Impact of OAS925 inoculation on alcohol and salt stress tolerance of *B. distachyon* plants. (A) photographs of OAS925 inoculated and uninoculated plant seedlings exposed to salt and alcohols, scale bar = 1 cm, (B) primary root length and (C) number of lateral roots. The error bars on the graphs represent the mean standard error of four biological replicates and colored dots represent the observed values of each replicate.

These findings confirm the phytotoxicity of short-chain alcohols, which is consistent with reports showing that such contaminants can damage plant cell membranes, inhibit germination, and alter nutrient uptake, ultimately reducing plant growth and biomass [2,21,22]. Contamination and exposure to even very low levels could reduce crop productivity drastically in agricultural fields. There are several reports of negative impact of other toxic solvents and hydrocarbon contamination on plants root growth [23–25]. The presence of these plant beneficial rhizosphere bacteria such as OAS925 can help these plants to survive and grow up to an extent since their tolerance levels are much higher than even the hardy grasses like *Brachypodium* as observed in this study. *Paraburkholderia* is one of the most stress tolerant bacterial genus present in the rhizosphere and endosphere of several plants [26–28]. Some of the commonly known plant protection mechanisms by *Paraburkholderia* are biofilm formation around the roots, production of phytohormones, increasing the nutrient availability and elevating the antioxidant levels to mitigate oxidative stress [28,29]. However, for further understanding of these mechanisms, it is first important that we understand how these bacteria are able to cope up in such stress conditions themselves in order to protect the plants.

### 3.2. RB-TnSeq identifies genes important for tolerance to short-chain alcohols in OAS925

To identify the genetic basis of alcohol and salt tolerance in OAS925, we performed a genome-wide fitness analysis using RB-TnSeq. Figure 3A presents a heatmap of gene fitness scores for loci exhibiting a strong fitness impact (|fitness| > 2.0 and |t| ≥ 4.0) in at least one experimental condition. Figure 3B shows a PCA (principal component analysis) plot derived from the fitness profiles of all significant genes, illustrating the global relationships between the cellular responses to each treatment. OAS925 was treated with a predetermined >50% inhibitory dose of these solvents (52 mM for isoprenol, 23 mM for prenol, 428 mM for ethanol, 349 mM for 2-propanol, 216 mM for propanol and 200 mM for salt. Overall, the fitness values of genes for all the alcohols had a high correlation with each other (except prenol) and were found clustered together while those for prenol and salt were closer to the control (glucose) and were clustered away from other conditions (Fig. S1). Several genes were observed to be significantly different in fitness (|fitness| under stress > 1.0 & |fitness| of control < 1.0, |t| ≥ 4.0) in the alcohol treatments compared to the control conditions (Table S1, Fig. 3A). Genes with significant fitness scores under individual stressors (compared to control) are mentioned separately in Table S2-S7. Clustering of these genes (|fitness| ≥ 2.0) based on the stress conditions showed similarity between ethanol and propanol and between isoprenol and 2-propanol (Fig. 3A). PCA of all the genes with significant fitness differences (|fitness| > 1.0 for stress and <1.0 for control, |t| ≥ 4.0), in all the treatments showed an overall high variance of ∼72% (Fig. 3B). Ethanol and propanol again were clustered in one group while isoprenol, 2-propanol and salt in the other and the controls (glu) and prenol were separately clustered away from others. Additionally, in all the alcohols except prenol, the majority of these genes showed fitness defects (negative fitness score) under their absence. However, in prenol, an opposite trend was observed with several of these genes having a fitness advantage (positive fitness score) in OAS925 and were clustered away from other alcohols (Fig. 3A-C).

**Fig. 3.**
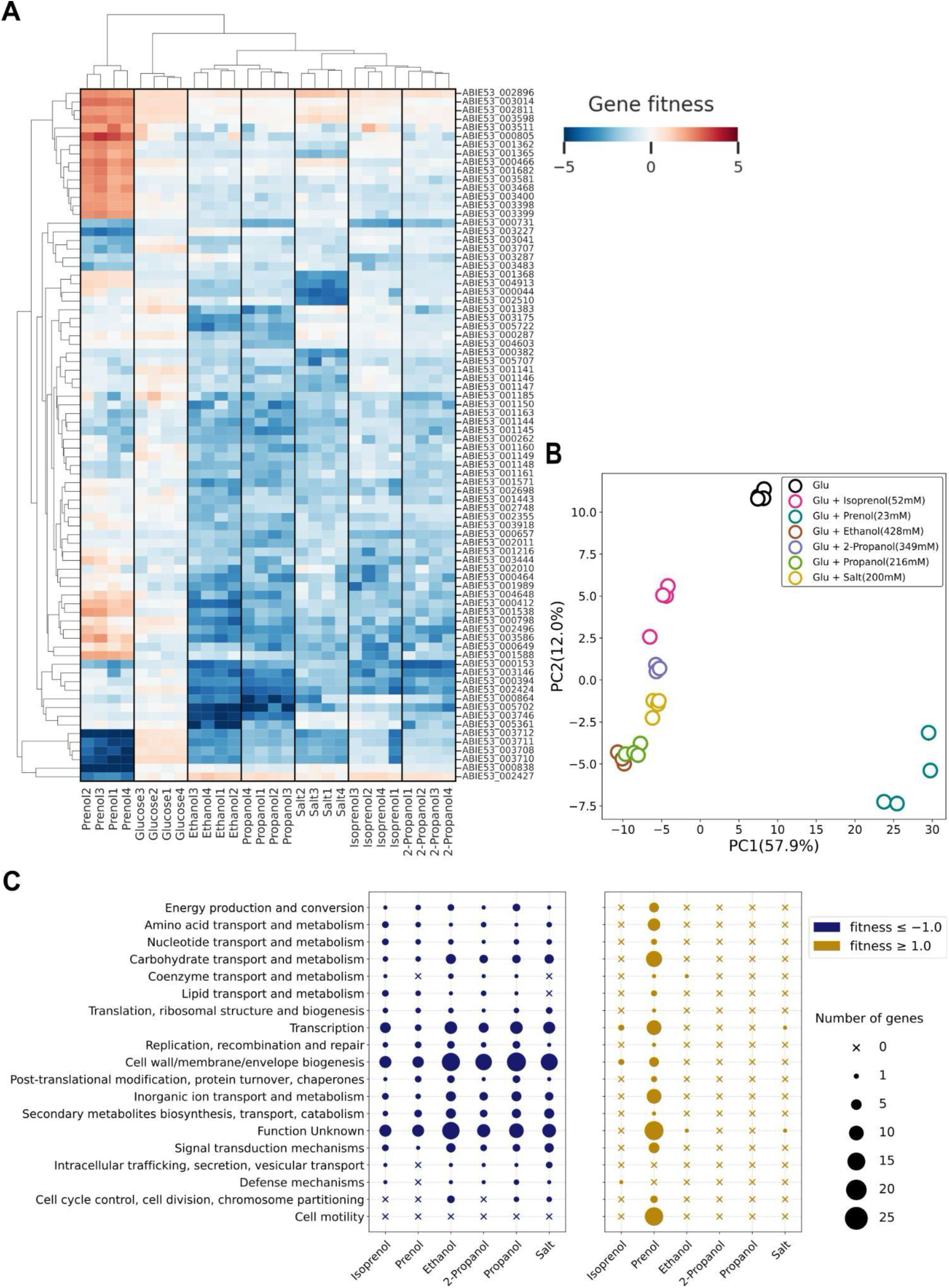
An overview of RB-TnSeq fitness data for OAS925 mutant library under alcohol and salt stress. A heatmap to the genes of OAS925 that had a significant fitness impact (|t| ≥ 4) in the presence of the salt and alcohols used in the study, only genes with |fitness| ≤ 2 are shown, for the list of all genes having a significant fitness impact ((|fitness| ≥ 1.0, |t| ≥ 4), see supplementary file (Table S1). The conditions were tested in quadruplicates (n = 4) and the data from each are numbered (1, 2, 3 & 4) (A). Principal Component Analysis (PCA) plot showing the clustering of all the significant genes based on the stress conditions (B). Number of OAS925 genes per clusters of orthologous groups (COG) category with significant fitness impact (|fitness| ≥ 1.0, |t| ≥ 4), involved in tolerance to alcohol/salt stress (C).

We predicted functions of all the genes with significant fitness scores based on clusters of orthologous groups (COG) of protein annotations, which was obtained using the eggNOG-mapper v2 tool [30]. Review of the COG categories showed that the majority of these genes were related to cell wall envelope/membrane biosynthesis, transport and metabolism, signaling systems, transcription/regulation, energy production and conversion, DNA repair, cell motility and cell division (Fig. 3C). The toxicity impacts of organic solvents including alcohols generally include increased cell membrane fluidity, altered regulation of internal pH, disruption of protein–lipid interactions, and decreased energy generation by the inhibition of transport of vital nutrients [31,32]. The significant fitness impacts on the genes that belong to such COG categories suggest that these alcohols may have similar toxicity mechanisms on OAS925. However, the cellular response to alcohol toxicity could also be dominated by the physicochemical properties of the alcohol rather than by the specific tolerance mechanisms in bacteria [33]. For instance, our study revealed a clear gene fitness clustering based on the branching of these alcohols i.e. straight-chain alcohols (ethanol and propanol) formed a distinct cluster from branched-chain alcohols (isoprenol and 2-propanol) with an exception of prenol which was clustered separately from other solvents. This suggested that bacteria exposed to solvents with similar chemical structure could have similar toxicity levels and tolerance mechanisms. A recent study by Arsov et al (2024), reported that bacterial strains were more tolerant to the toxicity from 2-butanol compared to 1-butanol and the transcriptomics data of *E. coli* ATCC25922 revealed reduced expression of genes for efflux pumps, chaperones, flagellar apparatus and enhanced expression of membrane and electron transporters compared to that in 1-butanol. Arsov et al discuss that the major reason for this difference in tolerance was the higher hydrophobicity of 1-butanol due to its unbranched structure [34]. We observed an overall difference between the fitness defect levels for several genes with lower fitness impacts for branched than unbranched alcohols (Fig. 3A). This difference was more pronounced in some of the genes for transporters (ABIE53_001144, ABIE53_001145, ABIE53_003710, ABIE53_003712), signaling system (ABIE53_005702) and chaperones (ABIE53_000864). It is likely that the differences in the hydrophobicity of these alcohols could be the reason for different tolerance mechanisms or the intensity of fitness impact. Solvents with greater hydrophobicities or partition coefficients tend to have higher membrane solubility, leading to disruption of lipid membranes in bacterial cells [35]. This could enhance the toxicity levels and could severely alter cell morphology [36,37].

*Paraburkholderia* is generally observed to be tolerant to salt stress and studies suggest its role in improving the salt tolerance in plants [38] [39]. Interestingly, the genes showing significant fitness impacts in the presence of salt made a separate cluster but were very close to other alcohols (except prenol) when all the significant genes were taken into account (Fig. 3B). This implies that some of these genes were similar fitness impacts to those found in the alcohols with few others that were specific to the salt stress tolerance. For instance, gene ABIE53_001368 which encodes for trehalose-phosphate synthase, is a crucial enzyme for synthesizing trehalose, a compatible solute which gets accumulated in bacterial cells in response to salinity stress [40].

Another interesting observation from the RB-TnSeq data was that the genes with significant fitness scores for prenol stress clustered away from all other conditions (Fig. 3A & 3B). In general prenol provided different trends relative to the overall trends observed in other solvents and possible rationales for this observation are discussed. One of the reasons could be the higher toxicity of prenol compared to other alcohols. Consistent with this, the toxicity level (∼50% lethal concentration) of prenol for OAS925 was also greater, with an MIC of ∼20mM relative to the other alcohols, ∼50mM (for isoprenol) to ∼420mM (for ethanol). Thus, it is possible that the bacteria need additional mechanisms to tolerate the extreme toxicity of prenol. Additionally, mutants in several genes such as those related to ABC transporters (ABIE53_003398-ABIE53_3400), response regulators (ABIE53_001362, ABIE53_001365) and cell division (ABIE53_003598, ABIE53_001538) exhibited a notable fitness advantage (fitness ≥ 1.0) in OAS925 when exposed to prenol, a contrast to their fitness defects observed with other alcohols (Fig. 3A, 3C). For example, absence of genes such as ABIE53_003598 (encoding cell division protein ZapD) and ABIE53_001538 (encoding Cytoplasmic axial filament protein CafA) confer a fitness advantage to OAS925 in the presence of prenol (Fig. 3A). These genes play a significant role in the cell division process under normal conditions, however, their absence may lead to change in cell shape and morphology which could be one of the survival mechanisms by bacteria under extreme stress conditions as observed in few studies [41–43].

Interestingly, this observed response to prenol may be unique to OAS925, given that in model organisms such as *E. coli*, prenol caused less growth inhibition compared to isoprenol [13]. Future studies could be focused on a detailed investigation to understand these differences.

#### 3.2.1 Genes for membrane-transporters important for bacterial fitness under alcohol stress

Among the genes with the most significant fitness defects were those involved in membrane transport. Figure 4A shows a heatmap of genes annotated encoding transporters that showed a strong fitness impact (|average fitness| ≥ 1.5, |t| ≥ 4) under one or more stress conditions. A set of genes namely ABIE53_003708, ABIE53_003710, ABIE53_003711, ABIE53_003712 that are homologous to *mlaC*, *mlaD*, *mlaE* and *mlaF* respectively, were important for fitness) under all the stressors tested in the study (Fig. 4A). These genes belong to the Mla (membrane lipid asymmetry) operon which is required for the transfer of lipid between the inner and outer membrane of bacteria and known to play a significant role in bacterial tolerance to various environmental stresses [44–46]. A schematic model of the Mla system’s proposed role in maintaining outer membrane lipid asymmetry in OAS925 is shown in Figure 4B. It is essential to maintain the lipid asymmetry in the outer membrane which acts as a barrier against several environmental stresses including those from exposure to toxic chemicals and solvents. For instance, the Mla pathway in *Burkholderia* has previously shown to be required for intrinsic resistance to extracellular stresses such as antibiotics and mutants lacking functional mla genes exhibited defective cell permeability and thus increased sensitivity to such stresses [46].

**Fig. 4.**
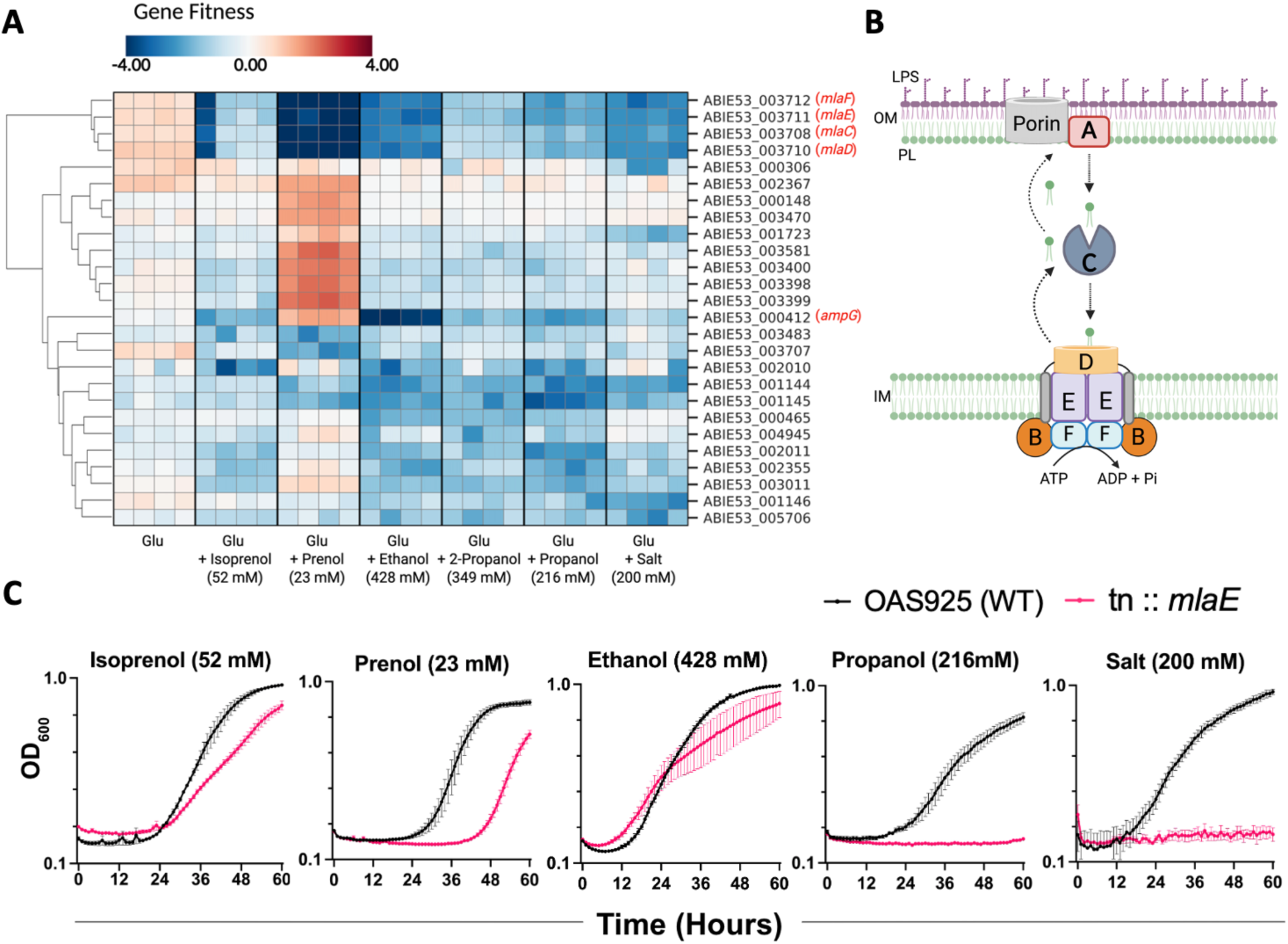
A heatmap for all the genes related to the transport and efflux systems that had a significant fitness impact (|average fitness| > 1.5, |t| ≥ 4) in the presence of the salt and/or alcohols used in the study (A). A schematic representation of the OAS925 mla system (B). Growth of WT OAS925 and a mutant in *mlaE* under the alcohols and salt stress conditions (C).

In addition to maintaining the membrane integrity, the Mla operon is also involved in biofilm formation, outer membrane vesicle formation, cell morphology and survival in several species [47,48]. Biofilm formation in the plant root environment could also be a major mechanism for providing the tolerance to these alcohols and solvents by providing an additional physical barrier against diffusion of these organic solvents and also in protection from osmotic stress caused by salt [49]. In a recent study, it was found that by enhancing rhizosphere colonization and biofilm formation, *Bacillus sp.* Za significantly reduced the toxicity of herbicide diphenyl ether to maize due to their enhanced herbicide degrading efficiency [50]. In our study, it is not clear whether OAS925 is capable of degrading these compounds, however, biofilm formation could still help the plants by decreased diffusion of these compounds and prevent the direct contact with the roots. As such we tested an individual *mlaE* mutant for the effect on growth under similar conditions to validate the results from RB-TnSeq and is presented in the Figure 4C. Relative to WT OAS925, an increase in lag phase and reduced growth was observed under all the conditions tested including some alcohols as well as salt validating the growth defect caused by the mutation in a single gene of the *mla* operon (Fig. 4C).

In addition to the *mla* operon, we also observed some other genes related to efflux systems that showed significant fitness defects in the presence of these alcohols. For instance, ABIE53_002355 which belongs to the Resistance-Nodulation-Division (RND) efflux system, outer membrane lipoprotein, NodT family had a significant fitness defect in the alcohol toxicity specifically isoprenol, ethanol and propanol (average fitness ≤ -1.5, |t| ≥ 4) (Fig. 4A). RND family transporters are mediators of multi-drug resistance in gram-negative bacteria and are mainly responsible for catalyzing the active efflux of many antibiotics and organic solvents [51]. RND transporter protein complex comprises cytoplasmic membrane transporter protein, a periplasmic-exposed membrane adaptor protein, and an outer-membrane channel protein. In addition to providing tolerance from toxic solvents to bacteria, these transporter complexes have also shown previously to play an important role in the interaction with plants by involving in initiation of colonization as well as in the survival of bacteria within plant tissue [52,53]. RND efflux proteins were found to be upregulated in *Pseudomonas putida* in the maize roots showing that they are also involved in the detoxification of inhibitory compounds present in plant root associated environments [54]. Thus, one mechanism involved in preventing plant roots from direct contact with potentially plant toxic alcohols could be by keeping them away via these efflux transporters, however, this needs further investigation.

Gene ABIE53_000412, encoding for an MFS transporter protein AmpG permease, was also required for fitness in the presence of alcohols, specially ethanol and propanol (Fig. 4A). In several gram-negative bacteria, it is involved in the transport of cell wall fragments required for peptidoglycan recycling [55,56]. In *E. coli*, it was found that AmpG permeases are involved in the production of beta-lactamases as well as biofilm formation as a resistance mechanism for beta-lactam antibiotics that interferes with the cell wall cross-linking [56]. There are limited reports of its role in *Burkholderia* or *Paraburkholderia* [57], it is, however, possible that these solvents may also interfere in the cell wall cross-linking activating AmpG in the similar manner and its absence could also detrimentally impact biofilm formation and survival of OAS95 under toxic conditions.

#### 3.2.2 Several genes related to signaling systems regulate the response to alcohol and salt stress

In addition to membrane transport systems, we found several genes encoding proteins related to two-component signaling systems (TCS) in OAS925 had significant fitness impact (|fitness| > 1.0, |t| > 4) under the majority of alcohol and salt stress conditions. These fitness candidates indicate that these solvents likely induce a very stressful environment for bacteria leading to the activation of these regulatory systems which play a vital role in protecting the bacteria against the alcohols and salt toxicity.

Gene ABIE53_002424 encodes for protein PrtR (Fig. 5A) and is predicted to be a part of TCS PrtI/PrtR where PrtI is an extracytoplasmic function (ECF) sigma factor and PrtR is the anti-sigma factor found in the membrane of many bacteria including those related to *Burkholderia* sp. [58–60]. The bacterial PrtR/PrtI system regulates the expressions for several phenotypes in response to environmental changes, such as production of extracellular protease (AprX/AprA), germination arrest factor (GAF) and lipases with few studies also suggesting its role cyclic lipopeptide (CLP) production which could be involved in cell motility [60–64]. We confirmed the fitness impact of some of these alcohols using an individual *prtR* mutant. Specifically, a longer lag phase and reduced growth was observed in the *prtR* mutant compared to WT on exposure to isoprenol, ethanol and propanol, validating the results obtained from RB-TnSeq analysis (Fig. 5B).

**Fig. 5.**
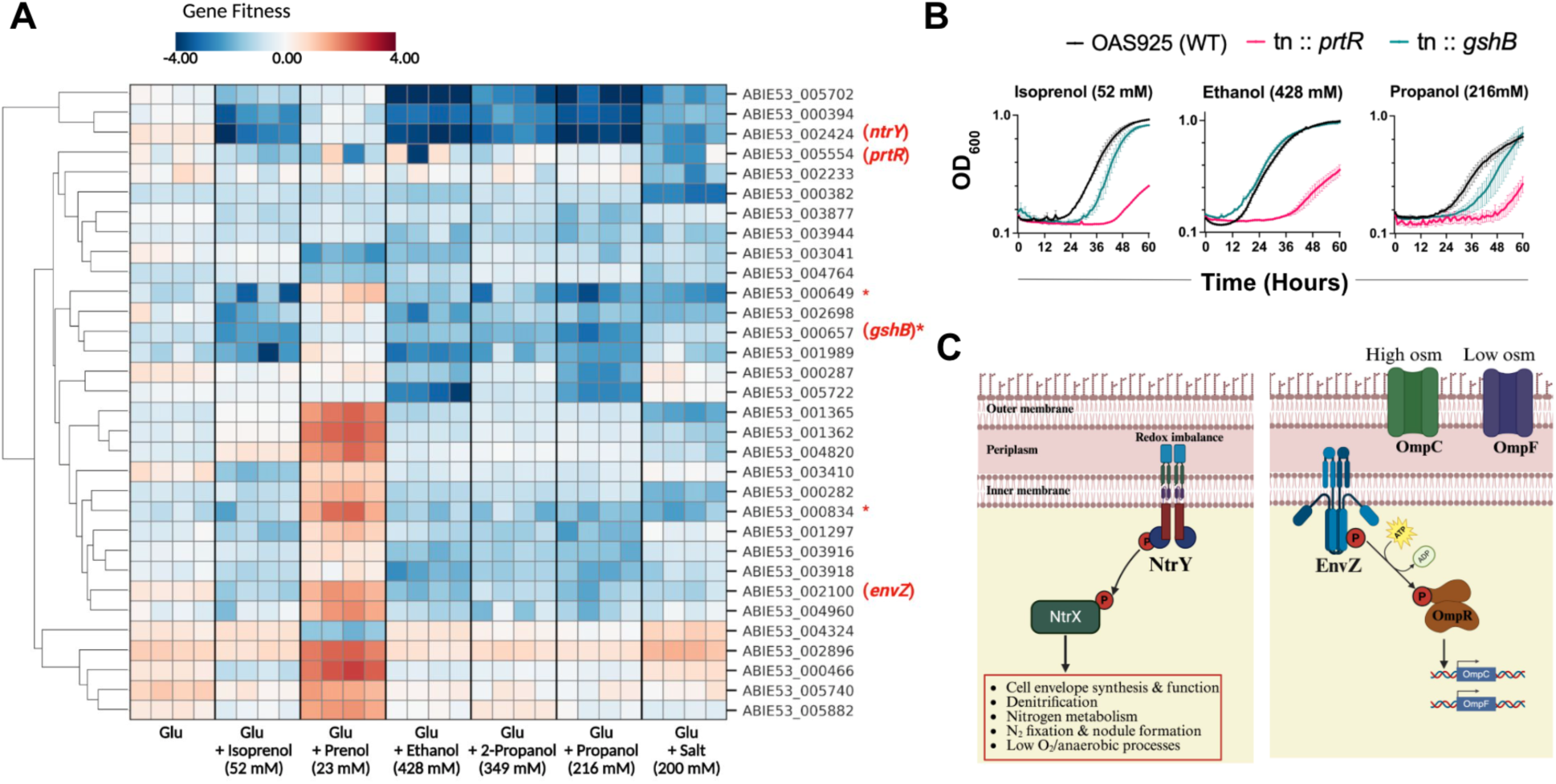
A fitness heatmap of genes related to signaling systems and oxidative stress (marked with *) that had a significant fitness impact in the presence of the salt and alcohols used in the study (A). Growth of WT OAS925 and individual mutants of *prtR* and *gshB* under the alcohols and salt stress conditions (B). Schematic representation of two component signaling systems NtrY/NtrX and EnvZ/OmpR involved in tolerance to alcohols and salt stress, osm : osmotic pressure (C).

Exposure to these solvents can cause changes in cellular osmolarity or redox imbalances, potentially triggering regulatory TCSs that control membrane protein function and maintain the integrity of the cell membrane and envelope structure [5,65–67]. Gene ABIE53_002100 encodes for EnvZ histidine kinase which is a component of the well-characterized bacterial EnvZ/OmpR system. This protein demonstrated a high fitness response to several alcohols and is known to modulate the response to osmotic stress (Fig. 5A, 5C) [68,69]. It is possible that these organic solvents cause desiccation or water exclusion similar to high osmotic pressure environments activating the EnvZ/OmpR system regulating the outer membrane porins (OmpC and OmpF) to prevent the solvents entering the cell. In a study by Y. Ma et al., EnvZ/OmpR was found upregulated in *Burkholderia sp.* to strengthen the negative regulation of OmpC and OmpF porins, effectively blocking phenol entering in the cells [70]. Another gene ABIE53_000394 encodes for NtrY protein, is a part of TCS NtrY/NtrX, where NtrY is the membrane associated histidine kinase and NtrX is the response regulator (Fig. 5A & 5C). This TCS is found in many alpha- and beta-proteobacteria and regulates several cellular processes like denitrification, survival in low oxygen/redox imbalances, cell envelope functions and nitrogen metabolism [71–73]. In *Rhodobacter sphaeroides*, loss of the NtrYX system impacted the levels of peptidoglycan precursors and lipopolysaccharides and altered cell envelope structure and functions resulting in increased outer membrane permeability [72,74]. Similarly, a transposon mutant of *ntrY* showed higher susceptibility in *Sinorhizobium meliloti* GR4 to salt stress with decreased motility, altered cell morphology and increased EPS production [75]. These reports suggest that the significant fitness defects observed in the absence of NtrY could be due to the alcohol/salt stress damaging the cell envelope leading to altered cell morphology and solvent permeability.

Taken together, these TCSs show an important role in regulating the bacterial defense against the toxicity from the introduced solvents, however, their role in directly or indirectly influencing the plant’s tolerance to these same stresses remains unclear. Studies have shown the role of these TCS in root attachment, colonization, EPS formation, biofilm formation and nutrient availability [62,75,76]. Regulation of such functions could have a very significant impact on plants tolerance to exogenously added stresses in the rhizosphere. For instance, Yang et al reported that the *prtR* mutants of *P. fluorescens* HC1-07 had impaired ability to persist in the rhizosphere of wheat due to their reduced surface motility and ability of biofilm formation compared to the wild-type strain [62]. In addition, the EnvZ/OmpR system has also been shown to influence bacterial motility, EPS formation and biofilm formation impacting its ability to colonize roots and establish symbiotic relationships with plants [76,77]. EnvZ/OmpR plays a crucial role in the transition from reversible to irreversible bacterial cell attachment during biofilm formation by inhibiting the synthesis of motile cell surface structures like flagella [78]. Apart from root attachment and biofilm formation, the TCS NtrX/NtrY is also involved in the regulation of nitrogen fixation process in several plant associated diazotrophs and symbionts such as *Azospirillum* and *Rhizobium* [73,79,80]. Plants could become nutrient deficient due to reduced amount of available nutrients in the presence of high salt or disruption of their nutrient absorption and uptake due to high concentration of toxic contaminants in soil [81,82]. While the nitrogen-fixing capabilities of OAS925 have not been studied extensively, it is worth noting that several *Paraburkholderia* species are known to fix nitrogen, either symbiotically or diazotrophically [29,83]. It can be speculated that these bacteria may help the plants under such conditions with fixing nitrogen and could be one of the plant protection mechanisms of OAS925 from alcohol and salt stress.

While our study implicates TCS in stress response of root associated bacteria, the precise molecular mechanisms that influence root attachment, biofilm formation, or beneficial functions like nitrogen availability require additional studies. A better understanding of these mechanisms is crucial, as they may significantly affect how root associated bacteria could protect the plants against the toxicity resulting from anthropogenically introduced solvents.

#### 3.2.3. Genes involved in protection from oxidative stress caused by alcohol and salt

Another gene found to be significantly involved in tolerance to these alcohols was for the ABIE53_000657 (*gshB*), encoding for glutathione synthase. Exposure to these solvents could induce oxidative stress in bacteria and plants potentially leading to the production of reactive oxygen species (ROS) such as hydrogen peroxide (H_2_O_2_) and the hydroxyl radical which can damage cellular components [84]. Bacteria can protect themselves from such oxidative damage by producing several antioxidant enzymes such as catalases, superoxide dismutases, and glutathione peroxidases, and antioxidants, such as the tripeptide glutathione γ-glutamyl-L-cysteinylglycine (GSH) synthesized by glutathione synthetases [85]. The redox-active sulfhydryl group of GSH protects cells from ROS by directly scavenging free radicals and acting as a cofactor for antioxidant enzymes such as glutathione peroxidases [85]. There are reports on the oxidative stress caused by the exposure to toxic solvents and chemicals as well as the role of antioxidant enzymes in the protection against the solvent toxicity in bacterial cells [86,87]. We further validated the results of RB-TnSeq by growing an individual mutant of *gshB* under similar conditions and observed a longer lag phase for the mutants compared to that of the WT OAS925 specifically under the toxicity of isoprenol and propanol (Fig. 5B).

Some studies also suggest that *Paraburkholderia* may protect plants from the oxidative stress induced by abiotic stresses by elevating the levels of antioxidant enzymes and antioxidant molecules in the plants [29]. However, more investigation is needed to confirm the direct role of antioxidant molecules produced in the bacterial cell in providing plant tolerance to the stress caused by these specific alcohols and salt.

### 3.3 OAS925 may protect plants from solvent toxicity by improved colonization and biofilm formation

RB-TnSeq data identified several genes, including those encoding membrane transporters or efflux systems, that play a role in helping the bacteria mitigate the toxicity of these solvents.. Many fitness targets genes observed in this study were also found to be related to root colonization, biofilm formation and bacterial motility, such as mla operon (section 3.2.1) and several TCS systems described previously (section 3.2.2). The *mla* operon has also shown to be involved in biofilm formation in many studies. In Figure 2 we show that OAS925 improved the root growth of plants exposed to these stressors. To determine if these significantly affected genes involved in root colonization or biofilm formation directly impact the bacterial ability to improve growth, we tested several mutants for their ability to colonize *Brachypodium* roots and form biofilms.

We observe a slight decrease in the root colonization efficiencies by the *prtR* mutant compared to WT under alcohol exposure. Importantly, the impact was more substantial particularly under isoprenol exposure to the roots (Fig. S2). For biofilm formation, we observed a similar trend of decrease in the biofilm formation on alcohol exposure in both WT and *mlaE* transposon mutant (Fig. 6A). A significant decrease in biofilm formation was observed for mutants on the roots of exposure to ethanol confirming the role of *mla* operon in biofilm formation on the roots.

**Fig 6.**
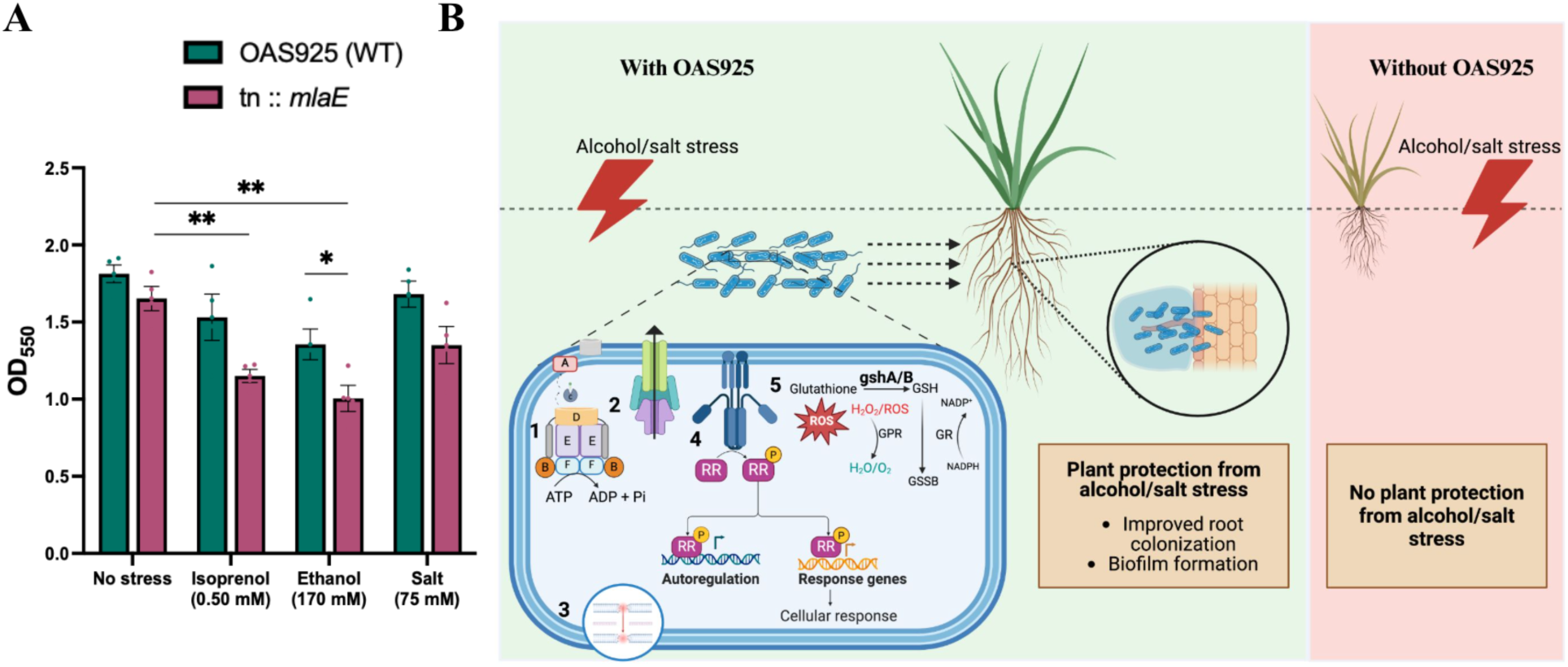
Quantification of biofilm formation on *B. distachyon* roots by *mlaE* transposon mutant compared to WT, the error bars on the graphs represent the mean standard error of four biological replicates, asterisk (*) represents significant differences with p <; 0.05 (*), p <; 0.005 (**) and no asterisk represents non-significant differences when t-test was performed between WT and mutant and between no-stress and stress conditions (A). Schematic representation of stress tolerance and plant protection mechanisms by OAS925, in the presence of alcohol/salt stress, OAS925 confers tolerance using several mechanisms such as transporters (1), efflux systems (2), maintaining membrane and cell wall integrity (3), two component signaling systems (4) and oxidative stress regulation (5), OAS925 also protects the plants by plausibly by improving colonization and biofilm formation on the roots (B).

These results suggest that biofilm formation and motility which impact the colonization of bacteria to roots could also be critical for as plant protection mechanisms involved in bacterial protection of plants from the solvent toxicity.

## 4. Conclusions

This study examines the detrimental impact of short-chain alcohols and salt, common biorefinery byproducts, on plant health, even at low concentrations. Most importantly, it reveals that the root-colonizing bacterium *P. graminis* OAS925 can significantly mitigate these negative effects, enhancing plant tolerance. Our findings point to four characteristics of bacterial resilience that directly support improvements to plant health: (1) the maintenance of outer membrane integrity and lipid asymmetry, governed by the *mla* operon; (2) the active removal of toxic solvents via RND-type efflux pumps; and (3) the coordinated regulation of stress responses through sophisticated signaling networks (PrtR/PrtI, NtrY/NtrX, EnvZ/OmpR), and (4) the mitigation of oxidative damage via glutathione synthesis (*gshB*). Ameliorating OAS925’s own stress response to enable growth may in turn enhance its competency to colonize roots for biofilm formation. The biofilm formation by OAS925 potentially acts as a synergistic protective barrier for plants against solvent toxicity. Together these findings offer a rudimentary mechanistic model for a beneficial plant-microbe interaction. Our work highlights the potential of plant-associated microbes like OAS925 in safeguarding agricultural systems from anthropogenic contamination and provides a foundation for future research into specific molecular interactions.

## Acknowledgements

The authors are thankful to John Vogel for providing seeds of *Brachypodium distachyon* to test for alcohol stress tolerance, and Guilherme M. V. de Siqueira for feedback on the manuscript. This material is based upon work done in Microbial Community Analysis & Functional Evaluation in Soils (m-CAFEs), a Science Focus Area led by Lawrence Berkeley National Laboratory supported by the U.S. Department of Energy, Office of Science, Biological and Environmental Research under Contract Number DE-AC02-05CH11231. J. C. M. has additional funding from the ARC Centre of Excellence in Plant for Space (CE230100015).

## Author contributions

S.P. and A.M. conceptualized the study. A.M.D. developed and provided the RB-TnSeq mutant library of *P. graminis* OAS925. S.P. performed the experiments in the lab and A.M.D. and V.T. did the sequencing and determined the gene fitness scores for all the experiments. V.T. isolated and provided the single transposon mutants to validate the results. J.C.M., T.E. and A.M. provided the supervision. A.M., J.C.M., and A.M.D acquired the funds for the project. S.P. drafted the initial manuscript. All authors have read, provided feedback, and approved the manuscript for publication.

## Supplementary Figures

**Fig. S1:**
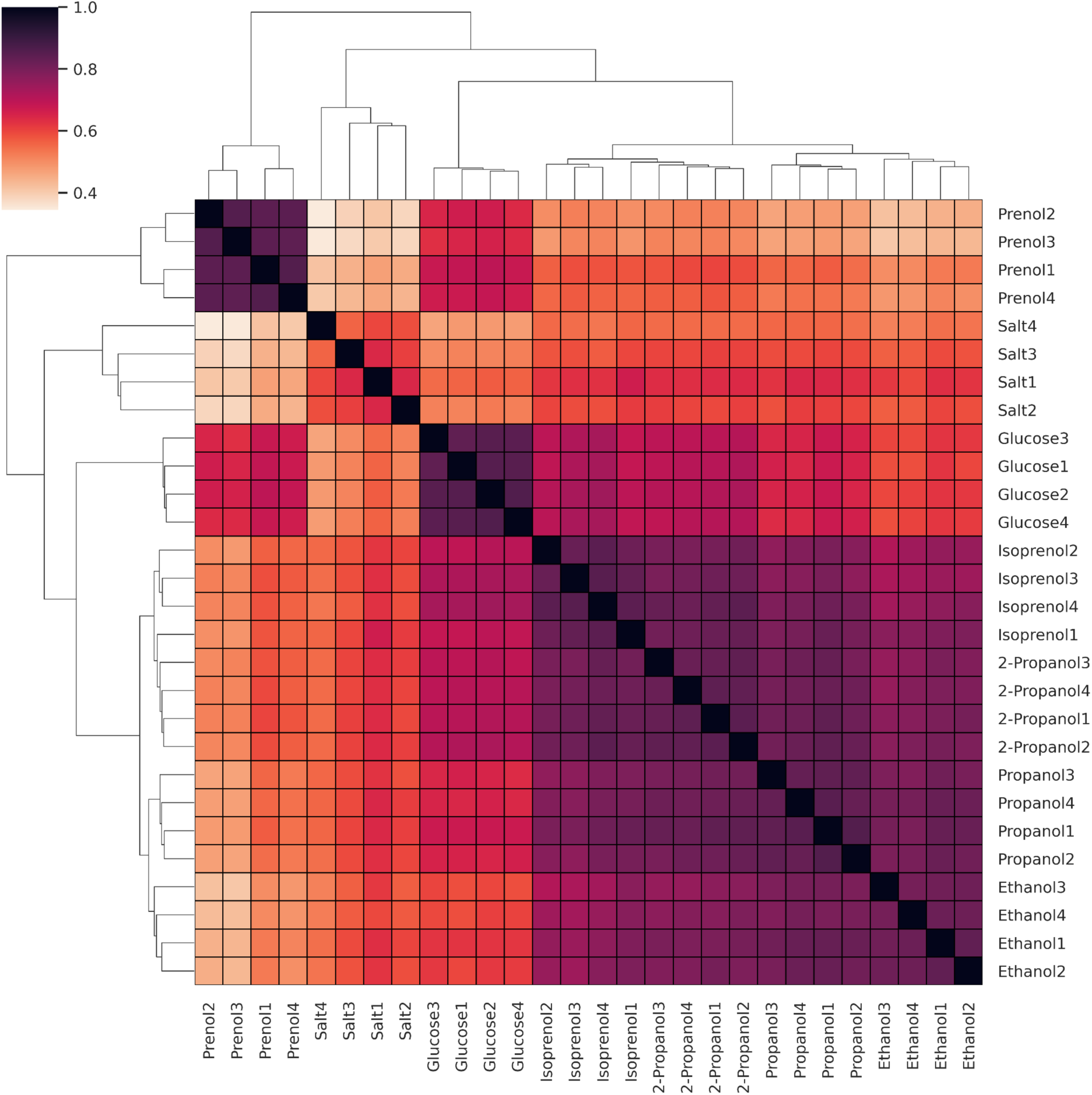
Cladogram correlation matrix of genome-wide fitness data of *P. graminis* OAS925 grown under alcohol and salt stress. The matrix shows pairwise comparisons of Pearson correlations of fitness data from OAS925 RB-TnSeq libraries grown in alcohols, salt as well as only glucose as control. The legend in top left shows Pearson correlation (r) between two conditions. The conditions were tested in quadruplicates (n =4) and the data from each are numbered (1,2,3 & 4).

**Fig. S2:**
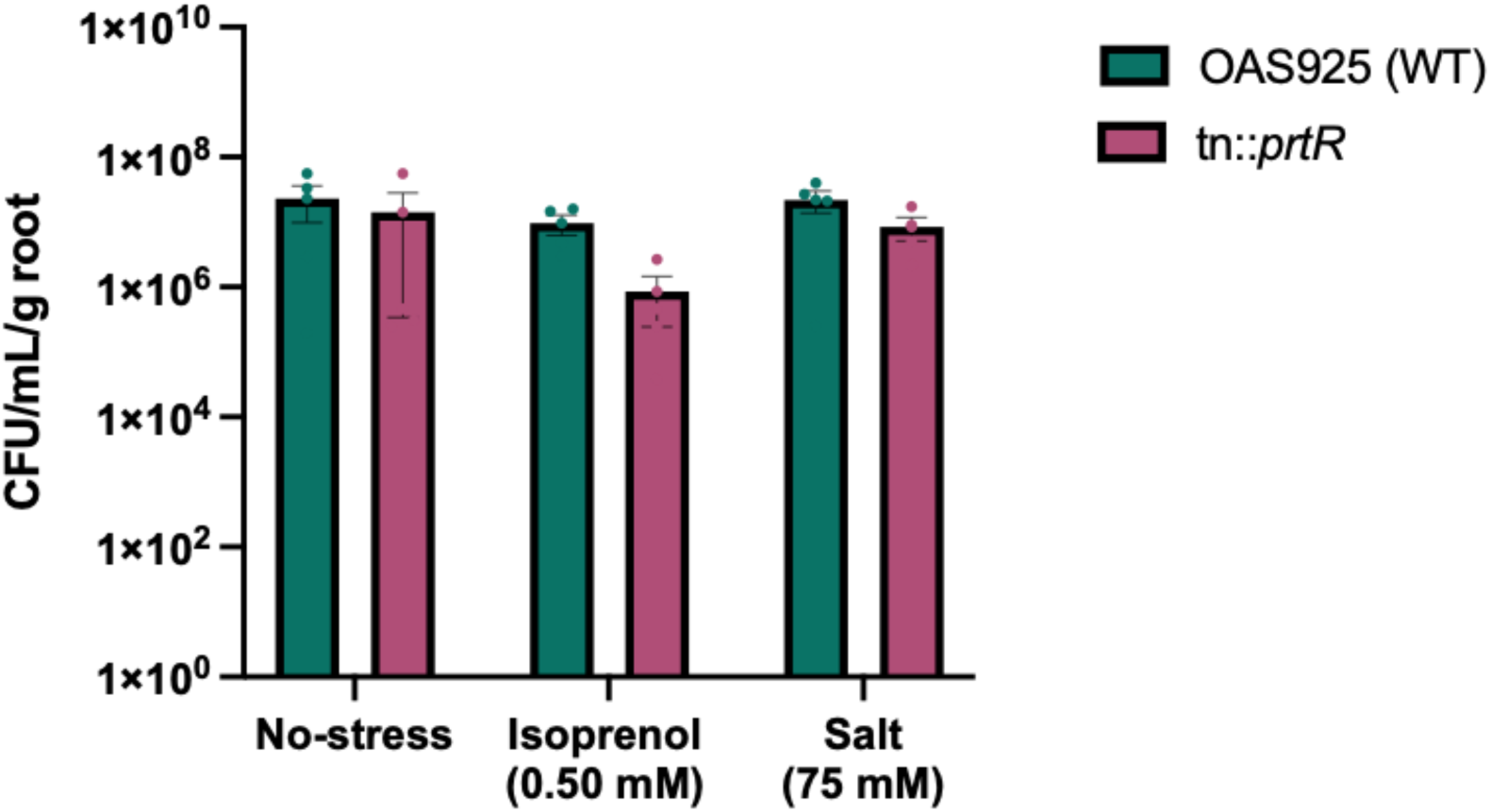
Plant root colonization test in the presence of isoprenol and salt inoculated with WT and *prtR* mutant of OAS925.

